# Direct observation of motor protein stepping in living cells using MINFLUX

**DOI:** 10.1101/2022.07.25.500391

**Authors:** Takahiro Deguchi, Malina K. Iwanski, Eva-Maria Schentarra, Christopher Heidebrecht, Lisa Schmidt, Jennifer Heck, Tobias Weihs, Sebastian Schnorrenberg, Philipp Hoess, Sheng Liu, Veronika Chevyreva, Kyung-Min Noh, Lukas C. Kapitein, Jonas Ries

## Abstract

Dynamic measurements of molecular machines can provide invaluable insights into their mechanism, but have been challenging in living cells. Here, we developed live-cell tracking of single fluorophores with nanometer spatial and millisecond temporal resolution in 2D and 3D using the recently introduced super-resolution technique MINFLUX. This allowed us to resolve the precise stepping motion of the motor protein kinesin-1 as it walks on microtubules in living cells. In addition, nanoscopic tracking of motors on microtubule of fixed cells enabled us to resolve their spatial organization with protofilament resolution. Our approach will enable future *in vivo* studies of motor protein kinetics in complex environments and super-resolution mapping of dense microtubule arrays, and pave the way towards monitoring functional conformational changes of protein machines at high spatiotemporal resolution in living systems.

---

Molecular machines drive many processes essential for life. For example, members of the myosin, kinesin, and dynein families are ATP-driven processive molecular motors that recognize the intrinsic structural polarity of cytoskeletal filaments to drive directed movement important for cellular processes like intracellular transport and cell division (*1*–*3*). Over the past decades, the microtubule plus-end directed motor kinesin-1 has served as a key model both for the understanding of motor dynamics and for the development of improved single-molecule methods, such as optical tweezers and single particle tracking (*4*–*7*). This has revealed that kinesin-1 moves in a hand-over-hand manner, in which each step along the microtubule encompasses a 16 nm displacement of the N-terminal motor domain, which leads to an 8 nm displacement of the C-terminal cargo binding domain (Fig. 1A). Despite the large success of existing single-molecule techniques in studying purified motors in well-controlled *in vitro* reconstitution experiments, performing such experiments in the living cell has proven challenging: large labels required for optical tweezers typically bind multiple motors in undefined ways and single fluorophores do not provide sufficient spatial and temporal resolution to resolve the fast stepping behavior of motors under physiological ATP concentrations. As a workaround, bright nanoparticles, taken up via endocytosis into transported vesicles, have been used as proxies to study motor dynamics in living cells (*8, 9*), but in such experiments the identity and copy numbers of the motors that drive transport are unknown. As such, the stepping dynamics of specific motors inside living cells has remained experimentally inaccessible.

**Figure 1:**
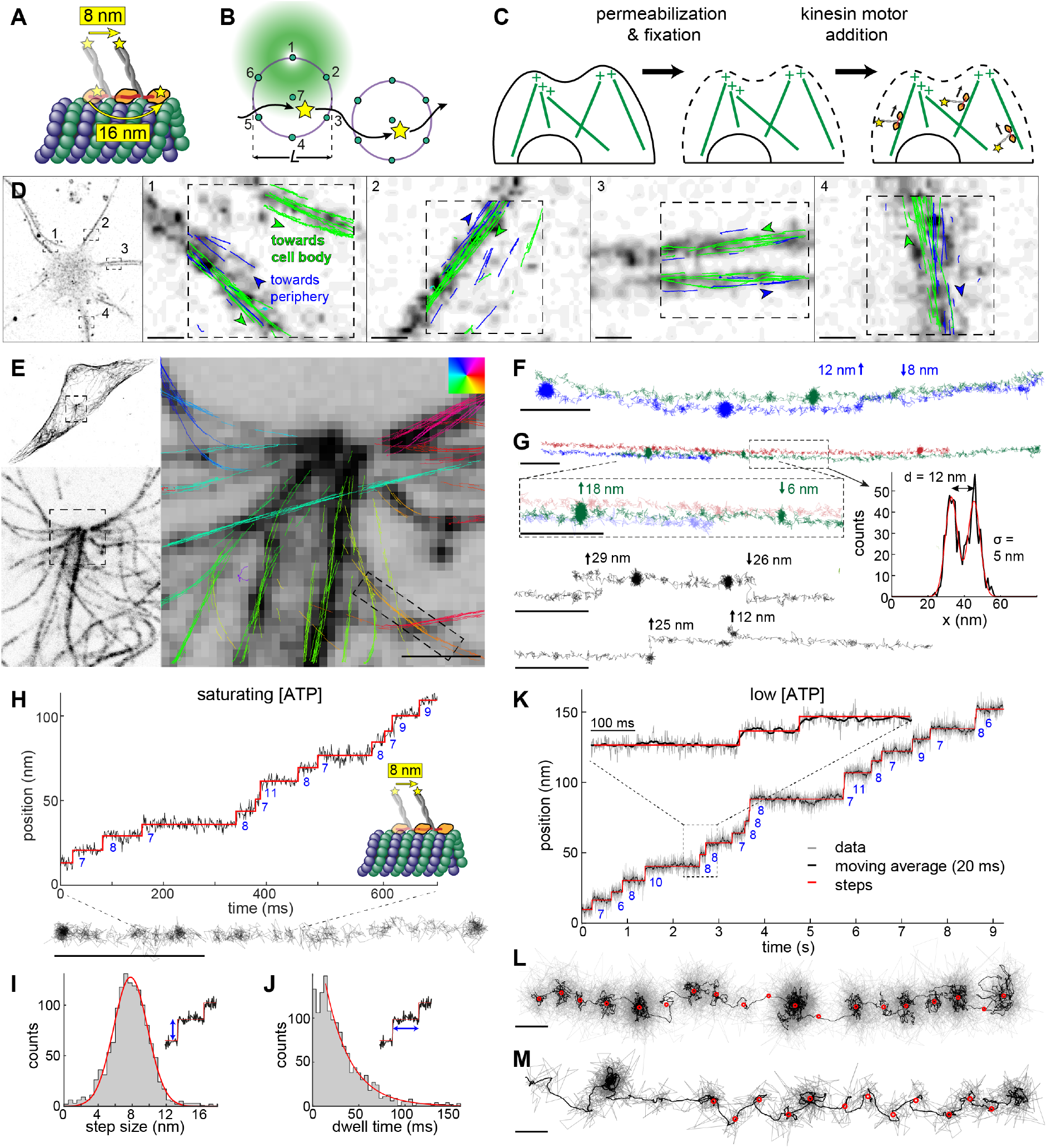
MINFLUX tracking of kinesin-1 in fixed cells. (**A**) **Kinesin-1** walks on microtubules (MTs) in a hand-over-hand manner. The apparent step size is 8 nm when the label is attached at the C-terminal tail domain and 16 nm when it is attached to the N-terminal motor domain. (**B**) **2D MINFLUX tracking** of a single molecule. A donut beam probes seven positions around a fluorophore to determine its location with nanometer precision. The scan pattern is iteratively centered on the fluorophore during tracking. (**C**) **Motor-PAINT** approach to track kinesin in fixed cells. Cells are first permeabilized to extract cell contents and then gently fixed to preserve MTs. Purified fluorescently labeled kinesins (*Dm*KHC (1-431)-SNAP-6xHis) are added and tracked as they walk towards the plus-ends of the MTs (movie S1). (**D** – **M**) **MINFLUX motor-PAINT in fixed cells**. (**D**) Confocal images of a neuron and overlaid kinesin trajectories in four neurites. Most of the neurites show bi-directional kinesin trajectories, i.e. towards (green) and away (blue) from the soma, as expected for dendrites. (**E**) Confocal microscopy images of GFP-α-tubulin in U2OS cells showing what appears to be a centrosome and overlaid kinesin trajectories with color-coded walking directions. (**F**) Tracks as indicated in (E) show side-stepping. (**G**) Tracks as close as 12 nm are clearly resolved and display kinesin switching laterally between neighboring MTs or protofilaments (movie S2). (**H**) Representative track and the corresponding time versus position plot at saturating ATP concentrations (>1 mM, here 6 mM), showing 8 nm walking steps. (**I**) Histogram of step size at saturating ATP concentrations from 7 experiments, 50 tracks, and 641 steps and Gaussian fit (7.8 nm ± 2.8 (std) ± 0.1 nm (sem); red line). (**J**) Dwell time histogram and fit to single exponential (*τ* = 27.0 ms; red line). (**K, L and movie S3**) A representative track at low ATP concentration (10 µM) and a corresponding time versus position, raw data (gray) and 20 ms running mean (black), clearly showing 8 nm walking steps. (**M and movie S4**) A representative track showing a zigzag trajectory, indicating an asymmetric arrangement of the label within a kinesin molecule (see fig. S2 for additional examples). Scale bars: 10 nm (L, M), 100 nm (F, G, H), 1 µm (D, E).

MINFLUX, a novel super-resolution microscopy technology (*10*), holds great potential to overcome these limitations. By efficiently utilizing the limited photon budget of single fluorophores, MINFLUX has reached unprecedented spatial resolution for imaging (*10*–*12*) and temporal resolution for fluorophore tracking (*13, 14*). In MINFLUX tracking, a donut-shaped excitation beam is scanned around a single fluorophore (Fig. 1B). From the intensities measured at specific positions, the coordinate of the fluorophore is calculated and the scanning pattern is re-centered on this position before the next iteration. Keeping the fluorophore close to the dark center of the beam results in a high localization precision and minimizes photobleaching. Nonetheless, whether this technique can enable high-resolution tracking of fast molecular motors in living cells has remained uncertain.

To establish MINFLUX tracking of molecular motors in living cells, we first optimized the workflow on single molecules and fluorescent beads *in vitro* to maximize precision and speed (fig. S1A-C) and achieved a localization precision of ≈ 2 nm with a sub-millisecond temporal resolution. We then optimized kinesin tracking using motor-PAINT (*15*). Here, cells are permeabilized and fixed, before fluorescently labeled motors are added that walk along microtubules towards the plus-end (Fig. 1C and movie S1). Motor-PAINT is less challenging than live-cell tracking as it allows us to precisely control the concentration of kinesin motors and, importantly, their speed via the ATP concentration. Using MINFLUX motor-PAINT, we were able to reconstruct cellular microtubules with a precision of ≈ 2 nm (Fig. 1D-E). Additionally, the directionality of kinesin reveals the orientation of the microtubules. Compared to standard motor-PAINT with a widefield microscope, the use of MINFLUX improved the localization precision 5-fold, the temporal resolution 50-fold, and the number of localizations per track by more than one order of magnitude. In neurons, this allowed us to better visualize the precise arrangement of anti-parallel microtubules inside dendrites compared to our earlier motor-PAINT study (Fig. 1D)(*15*). In human osteosarcoma (U2OS) cells, we could resolve individual trajectories in the crowded area around the centrosome with near-protofilament resolution (Fig. 1E, F). Tracks, just 12 nm apart, were easily resolvable and we regularly observed side-stepping between different protofilaments (Fig. 1F, G and movie S2), often after stalling, possibly to circumvent road blocks on the microtubules.

A closer inspection of individual tracks showed clusters of localizations that correspond to the 8 nm steps of the labeled C-terminus. Indeed, these steps become obvious when plotting the position of the motor along the microtubule over time (Fig. 1H). This allowed us to quantify precise step size and dwell time distributions under saturating (physiological) ATP concentrations (Fig. 1I, J). From 641 steps in 50 tracks, we measured a step size of 7.8 ± 2.8 (standard deviation, std) ± 0.1 (standard error of the mean, sem) nm. An exponential fit to the dwell time histogram resulted in a dwell time of *τ* = 27.0 ms. To investigate the stepping behavior in greater detail, we reduced the ATP concentration to slow down the motors (*16*). Under these conditions, we could measure hundreds to thousands of localizations per step (Fig. 1K, L and movie S3). Averaging over the coordinates allowed us to calculate the position of each step with sub-angstrom precision (sem). We note that currently, the accuracy of the measurements is not limited by the detected photons, but by the stability of the sample and microscope, as well as the offset of the label from the microtubule binding site. Interestingly, we noticed that more than 30% of the tracks displayed zigzag motion with every other step displaced perpendicular to the track center by ≈ 3-6 nm (Fig. 1M, fig. S2A and movie S4). This motion is likely due to an asymmetric positioning of the fluorophore with respect to the two motor domains imaged in a top view (fig. S2C) (*17*), demonstrating that MINFLUX can reveal intricate details of the conformational dynamics of individual motor proteins.

We next attempted MINFLUX tracking of kinesin in living cells. To this end, we expressed HaloTag-kinesin-1 (full-length) in U2OS cells and labeled at most a single motor domain per dimer by addition of the dye JF646 at very low concentrations. Individual tracks clearly revealed the 16 nm steps of the motor domains (Fig. 2A-D). On average, we found a step size of 15.7 ± 3.8 (std) ± 0.8 (sem) nm and a dwell time (exponential fit) of 31.8 ms (fig. S3A, B). We also observed tracks with frequent switching between microtubules, and, unlike in motor-PAINT, tracks without clear steps (fig. S4). The latter potentially stem from inactive kinesins that are attached to cargoes driven by other motors and are thus passively dragged along. However, most molecules were not bound to microtubules, but were unbound and displayed diffusive motions as expected for kinesin in an autoinhibited state (*18, 19*), limiting the number of tracks that we could acquire. To increase the throughput, we used the truncated kinesin-1 variant K560, in which cargo binding and auto-inhibition are removed, and treated cells with Taxol to increase the number of stabilized microtubules preferred by kinesin-1 (movie S5) (*20*). With these changes, we could readily observe multiple tracks in a single field of view (Fig. 2E-H and movie S6). This allowed us to measure precise *in vivo* step size and dwell time distributions from 2887 steps in 330 tracks (Fig. 2I, J). This revealed that, while the average step size of 16.2 ± 3.8 (std) ± 0.06 (sem) nm was similar to full-length kinesin, the dwell time of *τ* = 19.1 ms was much shorter, consistent with the higher speeds that we observed with K560 (table S1). To test whether our approach could be extended to much more complex and sensitive cell types, we examined kinesin-1 dynamics in the axons of live primary mouse cortical neurons (Fig. 2K-M), which critically depend on motor-driven transport. Also here, we could clearly quantify the 16 nm stepping dynamics of kinesin-1 (step size = 16.0 ± 4.0 (std) ± 0.5 (sem) nm, dwell time *τ* = 20.9 ms, fig. S3C, D), demonstrating that MINFLUX reveals conformational dynamics of individual motor proteins in complex cellular systems.

**Figure 2:**
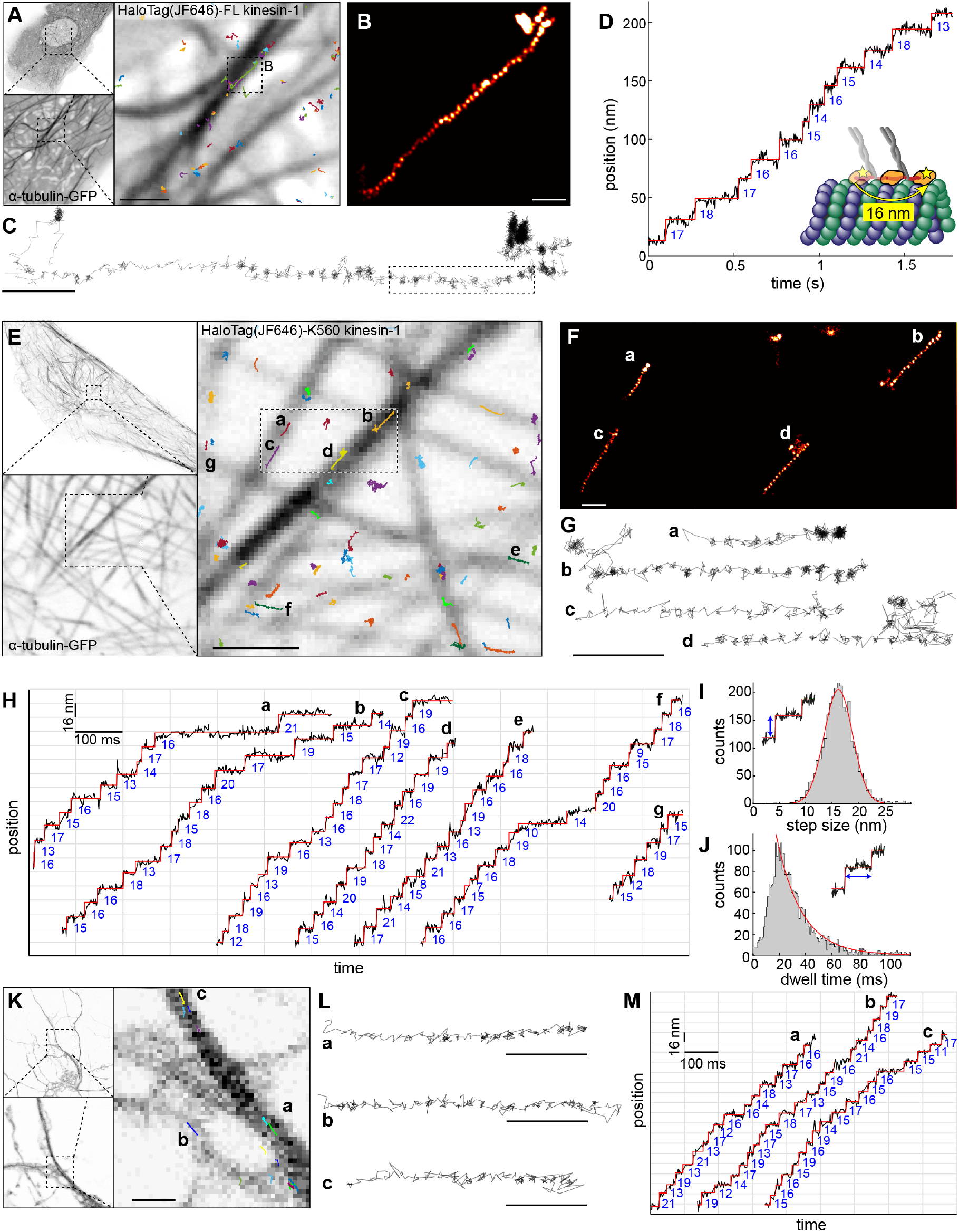
MINFLUX tracking of kinesin-1 in live cells. (**A** - **D**) **Tracking of full-length kinesin-1** labeled N-terminally with a HaloTag bound to JF646 in live U2OS cells. (**A**) Confocal images of GFP-α-tubulin in untreated live U2OS cells, and overlaid full-length human kinesin-1 trajectories. (**B**) A kinesin-1 track where the localizations are rendered as a super-resolution image, in the region indicated in (A). (**C**) A line plot connecting each localization. (**D**) Time versus position plot of the highlighted portion of the track in (C), showing walking steps of 16 nm. (**E** - **J**) **Tracking of truncated kinesin-1** (Halo-K560) in Taxol-treated live U2OS cells (movie S5). (**E**) Confocal images of GFP-α-tubulin and overlaid kinesin-1 tracks. The tracks indicated in (E) rendered as a super-resolution image (**F**), and line plots connecting each localization (G and movie S6), showing clear walking steps (localization precision: 2 nm; temporal resolution: 1 ms). (**H**) Time versus position plots of representative kinesin-1 tracks as indicated in (E), showing clear 16 nm stepwise movements. (**I**) Step size histogram (161 experiments, 330 tracks, and 2887 steps), fitted with a Gaussian function (*σ*= 16.2 ± 3.8 (std) ± 0.06 (sem) nm). (**J**) Dwell time histogram, fit with a single exponential (*τ* = 19.1 ms). (**K** - **M**) **Tracking of kinesin-1 (Halo- K560) in untreated live primary mouse cortical neurons**. (**K**) Confocal images of GFP-α-tubulin and overlaid kinesin-1 tracks. (**L**) Representative tracks as indicated in (K) as line plots and (**M**) time versus position plots, showing 16 nm stepwise movements(see fig. S3 step size and dwell time histograms). Scale bars: 100 nm (B, C, F, G, L), 1 µm (A, E, K).

Because most biological structures extend into three dimensions, only 3D tracking can capture the true dynamics and avoid projection artifacts that limit accuracy in 2D tracking. Unfortunately, standard single-particle tracking provides, at best, poor *z* resolution (*21*). MINFLUX has been used to image cellular structures in 3D (*11*), but for tracking it has so far remained limited to 2D. We therefore adapted MINFLUX for 3D tracking by scanning a 3D donut beam in 3D (Fig. 3A, B). We achieved a localization precision of 2.5 nm, 3.1 nm, and 3.9 nm in the *x,y,z* -directions, respectively (fig. S1D). When used with motor-PAINT, we could resolve many tracks on crossing microtubules (Fig. 3C) including jumps between microtubules (arrow in Fig. 3D). Importantly, we could also establish 3D tracking in live cells with a similar spatial and temporal resolution (4.2 nm and 2.8 ms), allowing us to resolve the 16 nm walking steps of kinesin in 3D (Fig. 3E, F, and movie S7). When we investigated these trajectories in the cross-sectional views, we found dynamic movements along the z-axis including side steps and vertical trajectories. Thus, MINFLUX tracking opens the possibility to quantify the precise 3D dynamics of molecular machines in living cells.

**Figure 3:**
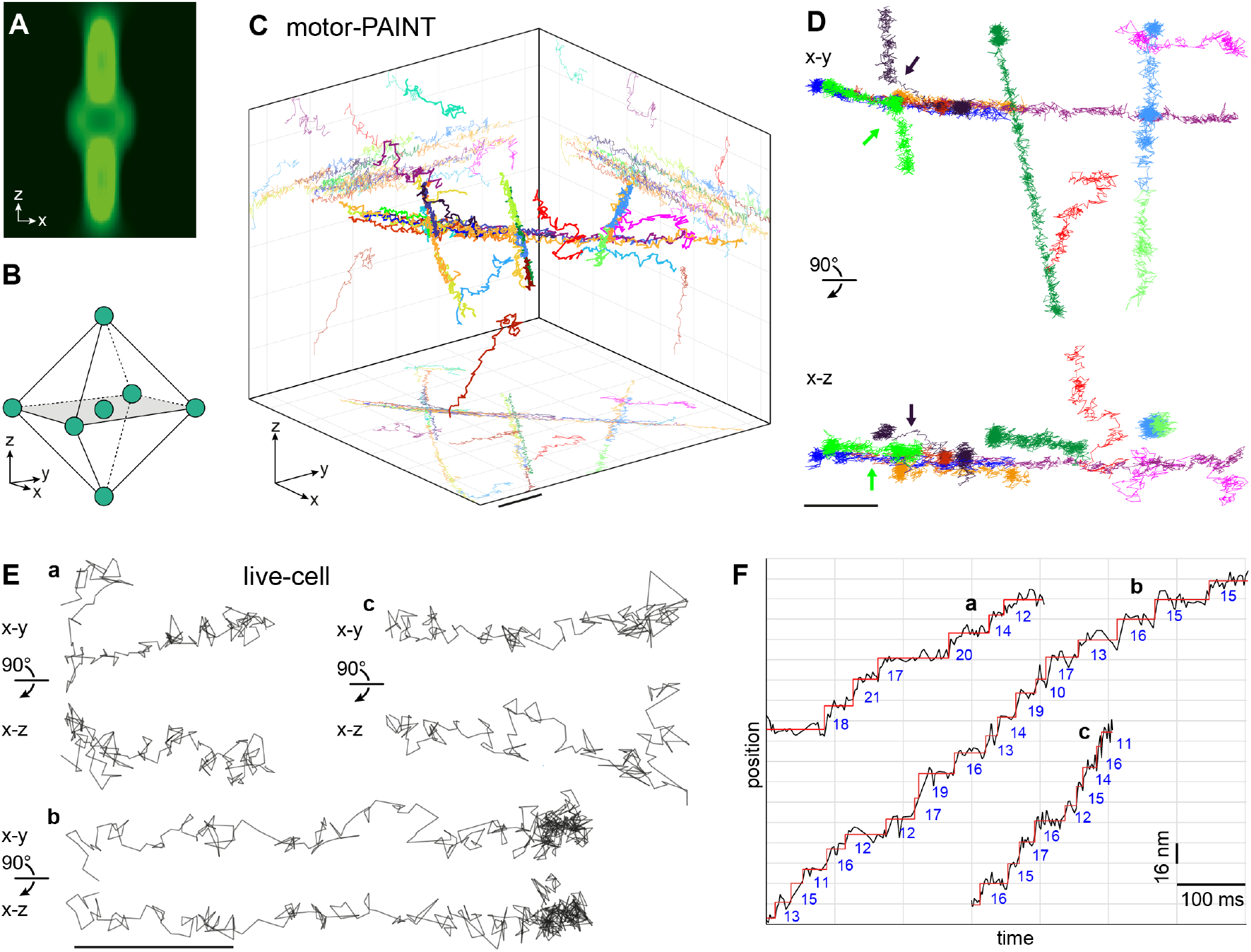
MINFLUX 3D tracking of kinesin-1. (**A, B**) **3D MINFLUX tracking**. (**A**) A 3D donut beam probes the intensity at (**B**) seven three-dimensionally distributed positions around a fluorophore. (**C, D**) **3D tracking with motor-PAINT** in fixed U2OS cells. (**C**) 3D rendering of kinesin-1 tracks at crossing MTs with a volume size of 1.2 µm × 1.2 µm × 1 µm. (**D**) Selected tracks from (C) in top and side views, including ascending and descending trajectories, and two trajectories in which motors switch MTs (arrows). (**E, F**) **3D tracking in live cells**. (**E**) Representative kinesin-1 tracks in live U2OS cells in top and side views, showing stepwise movements both in the x-y plane and along the z-axis (see fig. S6 confocal overview images and movie S7). (**F**) Position versus time plots of the tracks from (E), showing 16 nm steps. Scale bars: 100 nm (E), 200 nm (C, D).

We here established MINFLUX tracking of kinesin-1 with nanometer spatial and sub-millisecond temporal resolution. Using motor-PAINT, this approach allowed us to map precise positions and polarities of cellular microtubules with a resolution approaching the size of a tubulin dimer, one order of magnitude better than in standard motor-PAINT approaches. The combination of MINFLUX and motor-PAINT has great potential, as it not only allows probing kinesin dynamics in a controllable cellular system, but also enables mapping microtubules in extremely dense bundles such as the interpolar bundles in the mitotic spindle or the axonal microtubule array with unprecedented detail. In addition, the ability to discriminate motility along different protofilaments of the same microtubules might enable resolving defects or other alterations within the microtubule lattice. Importantly, we also demonstrated stepping of individual motors in live cells, which paves the way for studying how stepping behaviour is modulated by the presence or absence of different microtubule-associated proteins or cargo adaptors.

MINFLUX tracking is not limited to kinesin but can be used to study the precise motion of any protein in living cells with unprecedented spatio-temporal resolution and minimal perturbance thanks to single-fluorophore labels (see fig. S5 for tracking of Myosin-V). In the future, developing MINFLUX to simultaneously track two colors will enable monitoring the 3D relative positions of labeled protein domains with nanometer spatial and sub-millisecond temporal resolution. Such measurements of conformational changes of molecular machines in their native environment will provide important insights into their function and regulation.

## Acknowledgements

We thank Koki Watanabe for generating a full-length wild type Myosin-V construct, Abberior Instruments and specifically Roman Schmidt for MINFLUX technical supports, Angelica Pacheco, Ulf Matti, and Lucia Perez for their help in sample preparations, and the EMBL Imaging Centre for access to the MINFLUX instrument. JF646 Halo ligand was a kind gift of Luke Lavis, HHMI Janelia Research Campus. This work was supported by H2020 Marie Skłodowska-Curie Actions (RobMin, No. 101031734 to T.D.), EMBL ARISE fellowship (to S.L.), the European Research Council (grant no. ERC CoG-724489 to JR and CoG-819219 to L.C.K.) and the European Molecular Biology Laboratory (T.D., E.S., C.H., L.S., J.H., S.S., P.H., S.L, V.C., K.N., and J.R.).

## Author contributions

J.R. conceived the project. L.S., P.H., and V.C. designed and generated constructs. M.K.I. purified proteins for motor-PAINT. J.H. cultured neurons. T.D., M.K.I., E.S., C.H., L.S., J.H. prepared samples. T.D., T.W., and S.S. established MINFLUX tracking protocols. T.D. and S.L. acquired MINFLUX tracking data. E.S. and C.H. acquired fluorescence images. J.R. wrote analysis software. T.D., M.K.I., E.S., C.H., L.C.K., and J.R. analyzed the data. K.N., L.C.K., J.R. supervised the research. T.D., M.K.I, E.S., C.H., J.H., P.H., L.C.K., and J.R. wrote the manuscript with input from all authors.

## Methods

### Sample preparation

#### DNA constructs

To generate pHTN-HaloTag-KIF5B, full-length human KIF5B (1-963) was cloned from a HeLa cDNA library and inserted into the pHTN-HaloTag expression vector (both a kind gift from the Ellenberg Lab, EMBL Heidelberg) by AQUA cloning (*22*). The pHTN-HaloTag-KIF5B[1-560] construct (Halo-K560) was generated equivalently, except using different primers to obtain the truncated KIF5B gene (1-560). To generate pHTN-HaloTag-MYO5B[K1701A], full-length human MYO5 was cloned from the same cDNA library as above and inserted into the pHTN-HaloTag expression vector using the NEBuilder HiFi DNA Assembly kit (New England Biolabs, Ipswich, MA, USA). Subsequently, the point mutation K1701A was introduced by AQUA cloning. The sequences of all constructs were verified by Sanger sequencing (Eurofins, Ebersberg, Germany) or next generation sequencing (Plasmidsaurus, Eugene, OR, USA).

#### Protein purification

For motor-PAINT, *Dm*KHC (1-431)-SNAP-6xHis was purified from E. coli BL21 cells. Briefly, after transformation, bacteria were cultured until OD_600_ ≈ 0. 7 at 37°C. Cultures were cooled, after which protein expression was induced with 0.15 mM IPTG at 18°C overnight. Cells were then pelleted by centrifugation at 4500 × g, snap frozen in liquid nitrogen, and stored at −80°C until use. Cells were rapidly thawed at 37°C before being resuspended in chilled lysis buffer (50 mM sodium phosphate buffer supplemented with 5 mM MgCl2, 5 mM imidazole, 10% [v/v] glycerol, 300 mM NaCl, 0.5 mM ATP, and 1 × EDTA-free cOmplete protease inhibitor; pH 8.0). Bacteria were lysed by sonication (5 rounds of 30 s), supplemented with 2 mg/mL lysozyme, and then incubated on ice for 45 min. The lysate was clarified by centrifuging at 26000 × g for 30 min before being incubated with equilibrated cOmplete His-tag purification resin for 2 hrs. Beads were then pelleted and resuspended in 5 column volumes (CV) wash buffer (50 mM sodium phosphate buffer supplemented with 5 mM MgCl2, 5 mM imidazole, 10% [v/v] glycerol, 300 mM NaCl, and 0.5 mM ATP; pH 8.0) four times. Finally, the resin was transferred to a BioRad column. Once settled, the wash buffer was allowed to elute before adding 3 CV elution buffer (50 mM sodium phosphate buffer supplemented with 5 mM MgCl2, 300 mM imidazole, 10% glycerol, 300 mM NaCl, and 0.5 mM ATP; pH 8.0) to elute the protein. The eluent was collected, concentrated by spinning through a 3000 kDa MWCO filter, supplemented with 1 mM DTT, and 10% [w/v] sucrose before flash freezing in liquid nitrogen, and kept at −80°C. For labelling, protein was thawed and incubated with an additional 1 mM DTT for 30 min before adding 50 μM JF646-SNAP-tag ligand and incubating with rotation for 2 hours or overnight. Finally, excess dye was removed and protein was exchanged into wash buffer (low imidazole) supplemented with 2 mM DTT and 10% [w/v] sucrose by spinning through a 3000 kDa MWCO filter. Concentration was determined with a BSA standard gel. All steps from lysis onwards were performed at 4°C.

#### Cell culture

U2OS NUP96-SNAP cells (*23*) were cultured in DMEM growth medium with low glucose and without phenol red (Thermo Fisher Scientific, Waltham, MA, USA; Cat# 11880-028) supplemented with 1x MEM Non-essential amino acids (Thermo Fisher Scientific; Cat# 11140-035), 1x GlutaMax (Thermo Fisher Scientific; Cat# 35050-038), ZellShield (Minerva Biolabs, Berlin, Germany, Cat# 13-0050), and 10% [v/v] fetal calf serum (Thermo Fisher Scientific; Cat# 10270-106). Cells were grown in an incubator at 37°C, 5% CO2, and 100% humidity. High-precision 24 mm round glass coverslips (No. 1.5H; Marienfeld, Lauda-Königshofen, Germany; Cat# 117640) were cleaned by overnight incubation in methanol:hydrochlorid acid (50:50) stirring continuously, followed by subsequent washes with water until a neutral pH was achieved. Cleaned coverslips were then dried overnight under a laminar flow bench and cleaning was finalized by exposure to ultraviolet-radiation for 30 min. To utilize the sample stabilization system on the MINFLUX microscope, the prepared coverslips were coated with 0.01% poly-l-lysine (Sigma-Aldrich St. Louis, MO, USA; Cat# P4707) by applying 200 µL to each coverslip, incubating for 15 min, removing the solution, then applying 200 µL 200 nm gold nanoparticle solution (Nanopartz, Loveland, CO, U.S.A.) and incubating for 15 minutes under a running cell culture hood. Afterwards, the coverslips were rinsed twice with PBS and coating was finalized by UV radiation for 30 min. For imaging, 100k cells were seeded 3 days in advance on cleaned coverslips in a 6-well plate, each well containing 2 mL cell culture medium, to reach 50-70% confluency until the day of imaging. All experiments were performed with cells of different passage numbers.

#### Mouse primary cortex culture

Primary cortical neuron cultures were prepared from prenatal embryos of CD-1® IGS Mouse embryos (Charles River; Strain Code: 022; RRID: IMSR_CRL:022) at embryonic day 15 (E15). Animals were kept under standard SPF (specific pathogen free) conditions and sacrificed following routine and standard operating procedures of animal welfare with approval from the institutional animal care and use committee at EMBL. The embryonic cortex was dissected and collected in ice cold HBSS (containing Ca^2+^/Mg^2+^; Thermo Fisher Scientific; Cat# 14025092) from up to 15 mouse embryos. The tissue was washed thrice with ice cold HBSS (without Ca^2+^/Mg^2+^; Thermo Fisher Scientific; Cat# 14170112) and incubated with 0.25% Trypsin (Thermo Fisher Scientific; Cat# 15090046) in a water bath (37 °C) for 15 min. Afterwards, the tissue was washed again thrice with ice cold HBSS (without Ca^2+^/Mg^2+^) before dissociation/trituration in plating medium (Neurobasal; Thermo Fisher Scientific; Cat#: 12348017) containing 2% [v/v] B27 (Thermo Fisher Scientific; Cat#: 12587010), 1% [v/v] N2 (Thermo Fisher Scientific; Cat#: 17502048), 2% [v/v] GlutaMAX (Thermo Fisher Scientific; Cat#: 35050061), 1% [v/v] Pen/Strep (Thermo Fisher Scientific; Cat#: 15140122), 0.1 M NaPyr (Thermo Fisher Scientific; Cat#: 11360039), 10% [v/v] FBS (Thermo Fisher Scientific; Cat# 10270106), and 1 mg/mL DNAse I (Sigma Aldrich/Merck; Cat# 11284932001). The cell solution was passed through a 70 µm strainer and counted using Cellometer Auto T4 (Nexcelom Bioscience, Lawrence, MA, USA). 400,000 cells were plated on 24 mm coverslips pre-coated with 0.1 mg/mL Poly-D-Lysine (Sigma Aldrich; Cat# P0899), 2.5 μg/mL laminin (Sigma Aldrich; Cat#: 11243217001) and the gold beads. After 1 hour, a full medium change was performed using culturing medium (Neurobasal containing 2% [v/v] B27, 1% [v/v] N2, 2% [v/v] GlutaMAX, 1% [v/v] Pen/Strep, 0.1 M NaPyr). Cells were maintained at 37 °C in a humidified incubator with an atmosphere of 95% air and 5% CO_2_ until days in vitro (DIV) 8-10. A ⅓ to ½ medium change was performed twice a week.

#### Motor-PAINT

For motor-PAINT, U2OS cells or neurons were permeabilized for 1 minute in extraction buffer (BRB80: 80 mM K-PIPES, 1 mM MgCl2, 1 mM EGTA; pH 6.8, supplemented with 1M sucrose and 0.15% [v/v] TritonX-100) pre-warmed to 37°C. Pre-warmed fixation buffer (BRB80 supplemented with 2% [w/v] formaldehyde (FA) was added to this (i.e. final FA concentration of 1% [w/v]) and the solutions were mixed by gentle pipetting for 1 min. Subsequently the extraction-fixation-solution was exchanged for prewarmed wash buffer (1 µM paclitaxel in BRB80) and incubated for 1 min. The 1-minute wash step was repeated three more times, and staining solution (1 to 1000 dilution, BioTracker 488 Live Cell Microtubule Dye, Sigma-Aldrich, in BRB80) was added and incubated for at least 1 hr to enhance the microtubule staining (Fig. 1D, G, 3C, D). This solution was removed, and the chamber was washed one time for 1 min in the wash buffer (BRB80 supplemented with 1 µM Taxol). The staining solution was then exchanged for prewarmed imaging buffer (BRB80 supplemented with 583 µg/mL catalase, 42 µg/mL glucose oxidase, 1.7% [w/v] glucose, 1 mM DTT, 2 mM methyl viologen, 2 mM ascorbic acid, 1 µM Taxol, and 5 mM ATP) containing ≈ 0.5 nM kinesin.

#### Transfection of U2OS cells

One day after seeding, cells were transfected using lipofectamine 2000 (Thermo Fisher Scientific; Cat# 11668019). For each coverslip, 3 µL lipofectamine together with motor protein plasmid DNA (50 ng/coverslip), GFP-α-tubulin (100 ng/coverslip) (*24*), and salmon sperm DNA (850 ng/coverslip) were added to 100 µL Opti-MEM Reduced Serum Medium with GlutaMAX Supplement (Thermo Fisher Scientific; Cat# 51985026) and mixed well by vortexing. After a 10 min incubation, the cell culture medium was exchanged for 2 mL Opti-MEM and 100 µL of the transfection solution was added dropwise per coverslip. Cells were cultured in an incubator at 37 °C and 5 % CO_2_ overnight and the transfection medium was exchanged for cell culture medium the next day. Kinesin was co-transfected with GFP-α-tubulin (*24*) while MYO5 was co-transfected with GFP-Lifeact.

#### Transfection of mouse primary neurons

Primary mouse neurons were transfected at DIV2 or 3 using calcium phosphate precipitation. For one coverslip (Ø 24 mm), sterile double distilled H_2_O, (added up to 60 µL) was mixed with 6 µL of 2 M CaCl_2_, and 6 µg total endotoxin free plasmid DNA (4 µg HaloTag-KIF5B1-560 plasmid DNA + 2 µg of α-tubulin-GFP) isolated using the NucleoBond Xtra Midi Plus EF kit (MACHEREY-NAGEL GmbH & Co. KG, Düren, Germany; Cat# 740422.50). The mixture was incubated for 5 min at room temperature (RT). Next, 60 µL of 2× HBS buffer (274 mM NaCl, 9.5 mM KCl, 1.4 mM Na_2_HPO_4_, 15 mM HEPES, 15 mM glucose, pH 7.14) was added followed by vortexing briefly. The mixture was then incubated for 25 min at RT. Prior to transfection, conditioned culturing medium was removed from the cells (and stored) and replaced with Neurobasal (Thermo Fisher Scientific; Cat# 12348017) stored in the incubator at 37°C overnight. Subsequently, 120 µL of the transfection mixture was applied dropwise per coverslip followed by an incubation of 30-40 min at 37°C in the incubator until the precipitate was formed. The cells were washed two times with 1× HBS (pH 6.9) for 5 min in the incubator and finally once with Neurobasal (kept in the incubator overnight) for 5 min in the incubator. After the Neurobasal wash, the medium was replaced with the original conditioned culturing medium and cells were cultured until DIV8-10 for the live cell imaging experiment.

#### Live cell staining and microtubule stabilization

For live imaging of transfected cells, HaloTag live labeling of recombinant kinesin was performed with the JF646-HaloTag ligand (Janelia, Ashburn, VA, USA) on the morning of imaging. Cell culture medium was exchanged for 0.5 nM JF646 in PBS and incubated for 15 min at 37 °C and 5 % CO_2_ before changing back to culture medium. For experiments with Taxol for stabilization of microtubules, the coverslip of interest was incubated with 10 µM paclitaxel (Taxol, Sigma-Aldrich) one hour prior to imaging, by addition to the cell culture medium and the cells were kept in the same media throughout imaging.

#### Test sample preparation

For optimizing 2D tracking parameters, single Atto647N nanobody (20A_Atto647N, Biomers.net, Germany) deposited on a glass coverslip was used. We immobilized the dye by the following protocol. First, the coverslip surface was plasma-cleaned (Plasma Prep2; PIE Scientific, U.S.A.) for 5 min. Then we applied a solution of 2% APTES (Sigma-Aldrich, U.S.A.), 5% of water and 93% of Ethanol, incubated for 15 min at RT, washed twice with 100 % ethanol, washed twice with DI water, and placed the sample at 65 °C for 1.5 hrs. Then, a dye solution in DI water (1:100,000 dilution) was applied and the coverslip was incubated for 20 min at RT. After the incubation, the coverslip was washed with 100 mM NH_4_Cl three times, washed with DI water for three times, and was dried overnight. For optimizing 3D tracking, fluorescent beads (50 nm TetraSpeck microspheres fluorescent blue/green/orange/dark red, C47281, Thermofisher, U.S.A.) were used. The beads were immobilized on a glass coverslip by incubating in 100 µM MgCl2 in water for 10 min and then the coverslip was washed twice with PBS.

#### Sample mounting

Samples were mounted on cavity slides (Brand GmbH & Co KG, Wertheim, Germany, Cat# 475505) containing approximately 150 µL imaging buffer for motor-PAINT, cell culture medium containing 20 mM HEPES buffer (pH 7.25) for live cells, air for the 2D tracking test sample, and PBS for the 3D tracking test sample. Edges of the coverslip were sealed with dental addition-curing duplicating silicone (Picodent twinsil; Dental-Produktions-und Vertriebs-GmbH, Wipperfürth, Germany) and cured for 5 min at 37 °C.

### Microscopy

#### MINFLUX microscopy

A commercial MINFLUX microscope, Abberior MINFLUX (Abberior Instruments, Göttingen, Germany)(*14*), was used for all MINFLUX tracking experiments. In detail, we used a 100x oil immersion objective lens (UPL SAPO100XO/1.4, Olympus, Tokyo, Japan), 642 nm CW excitation laser, two avalanche photodiodes (SPCM-AQRH-13, Excelitas Technologies, Mississauga, Canada) with a detection range of 650 – 685 nm for the first detector and 685 – 760 nm for the second (detected photons were summed), and a pinhole size corresponding to 0.78 airy units. All the hardware was controlled by Abberior Imspector software (version 16.3.13924-m2112). The drift during the motor-PAINT measurements was minimized using the built-in stabilization system of the Abberior MINFLUX with typical drifts under 1 nm in xyz directions. For the sample stabilization, 200 nm gold nanoparticles (Nanopartz, Cat# A11-200-CIT-DIH-1-10) deposited on the coverslip surface were used as a positional reference.

#### Tracking parameters and sequences

We applied an iterative localization approach (*11*) with decreasing L size, a different number of collected photons, and a different dwell time in each iteration, as listed in the tables below. During the last iteration, the system continued localizing a molecule with the same parameters until its abort criteria (either losing the detection signal or exceeding the predefined central frequency ratio (CFR) value) was met. The CFR is a ratio of detection frequencies at the center and at the offset (position 7 vs other positions, Fig. 1A), and was introduced in previous work (*14*) to reject biased localizations due to multiple emitters close to the targeted coordinate patter (TCP).

The following two tables show the parameters used for 2D and 3D tracking, respectively.

| **2D tracking** | **1<sup>st</sup> iteration** | **2<sup>nd</sup> iteration** | **3<sup>rd</sup> iteration** | **4<sup>th</sup> iteration** |
| --- | --- | --- | --- | --- |
| **L size (nm)** | 284 | 302 | 151 | 75 |
| **TCP** | Hexagon | Hexagon | Hexagon | Hexagon |
| **Number of collected photons (counts)** | 40 | 20 | 10 | 100 |
| **Laser power multiplier \*** | 1 × | 1 × | 2 × | 2.5 × |
| **TCP dwell time (μs)** | 50 | 50 | 50 | 300 |
| **Pattern repeat** | 1 | 1 | 1 | 3 |
| **CFR** | -1.0 | 0.5 | -1.0 | 0.9 |
| **Background threshold (kHz)** | 15 | 30 | 30 | 40 |

\* This is a multiplier on the reference laser power to achieve an actual power used in each iteration.

| 3D tracking | 1<sup>st</sup> iteration | 2<sup>nd</sup> iteration | 3<sup>rd</sup> iteration | 4<sup>th</sup> iteration | 5<sup>th</sup> iteration |
| --- | --- | --- | --- | --- | --- |
| L size (nm) | 288 | 1440 | 288 | 151 | 100 |
| TCP | Hexagon | Zline** | Octahedron | Octahedron | Octahedron |
| Number of collected photon (counts) | 40 | 30 | 40 | 40 | 300 |
| Laser power multiplier* | 1 × | 1 × | 1 × | 2.5 × | 6.0 × |
| TCP dwell time (μs) | 200 | 200 | 200 | 200 | 300 |
| Pattern repeat | 1 | 1 | 1 | 1 | 1 |
| CFR | -1 | - 1 | 0.5 | - 1 | 1.0 |
| Background threshold (kHz) *** | 30 | 30 | 35 | 35 | 50 |

\* A multiplier on the reference laser power to achieve an actual power used in each iteration.
\*\* A TCP for an axial scan with two points above and below the focal plane.
\*\*\* In 3D tracking, an automatic background estimation feature was enabled. The actual background threshold value was the sum of the automatically estimated value and the value listed in the table.

Reference laser power values for each experiment, measured at the back aperture of the objective lens, are listed in the table below.

|  | Fig. 1D to F | Fig. 1 G to L | Fig. 2 A to M | Fig. 3 |
| --- | --- | --- | --- | --- |
| Reference laser power (μW) | 37 | 110 | 110 | 37 |

#### Imaging for parameter optimizations

For determining the tracking parameters, we first optimized a background threshold value in the following way. With the parameters listed in the table, we switched off the automatic background threshold function and set the background threshold to zero so that the system tracked the background at the last iteration. 5 kHz was added to the median value of the measured signals and was used as a threshold value for 2D tracking. By doing the same but with the automatic background estimation function on, we added 40 kHz for the 3D tracking. Then, we optimized the laser power (i.e. laser power factor) by tracking immobilized single Atto647N molecules and finding the power which gave a ≈150 kHz detection frequency. In addition, we tested different numbers of pattern repeats with a triangular and hexagonal TCP. Based on the overhead time for the pattern repeats, we set the repetition to 1 repeat per 100 μs, so it does not slow down our target temporal resolution (1 kHz localization rate) significantly but achieves sufficient averaging to counteract system instability. Finally, we optimized the number of photons to reach a ≈ 2 nm localization precision, which was over 100 photons per localization. For 3D tracking, we followed the same procedure but with a 40 nm fluorescent nanosphere in water instead of air.

#### Widefield microscopy

For a sample quality check to confirm kinesin motility in both motor-PAINT and live cells (movie S1, S5 for motor-PAINT and Halo-K560, respectively), we used a custom-built standard fluorescence microscope. The microscope details were described previously (*25*). Briefly, we used a 100× silicon oil immersion objective lens (UPlanSApo 100×/1.35 Silicon oil objective lens, Olympus, Japan), 638 nm excitation laser at 1.9 mW (iBeamMLE; Toptica, Munich, Germany), detection wavelength of 685/70 nm, and imaged on a sCMOS camera (Orca-Flash 4.0 V2, Hamamatsu Photonics, Japan) with a projected pixel size of 123 nm and an exposure time of 30 ms. The microscope was controlled by MicroManager 2.0.0 (*26*) with htSMLM (*27*) for image acquisitions.

### Data processing and analysis

All MINFLUX data was processed, analyzed, and rendered on a custom MATLAB software, SMAP (*28*). Motor-PAINT data was processed with the motor-PAINT plugin to obtain steps, averaged trajectories without long stops, and orientations of the tracks. Localization data was rendered into a super-resolution image as a convolution of localizations with a Gaussian function with σ = 3 nm (Fig. 2B and F). Steps in acquired tracks were analyzed in the ROI manager using the stepsMINFLUX plugin, whose step finding algorithm is based on a previous study (*29*). We chose specific regions within the tracks when necessary and/or corrected the found steps when the algorithm failed to correctly identify steps.

We estimated the experimental localization precision *σ* for each track by calculating the standard deviation of coordinate difference between consecutive localizations:

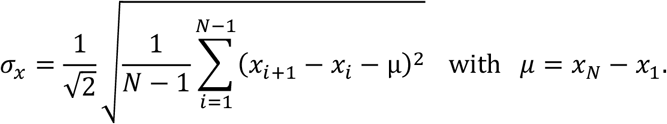

For static localizations with only random, uncorrelated noise, the variance of the coordinate differences is twofold higher than that of the coordinates themselves, leading to a 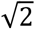 higher *σ* when calculated on the differences compared to when calculated on the coordinates directly. For mobile fluorophores, this equation over-estimates the noise. Correlated noise (e.g., drifts) on the other hand is underestimated.

For a statistical analysis of the found step sizes, we used Gaussian function fitting. For the dwell time analysis, a single exponential curve was fitted from the peak of the histogram.

## List of Supplementary Figures and Tables

**Fig. S1:**
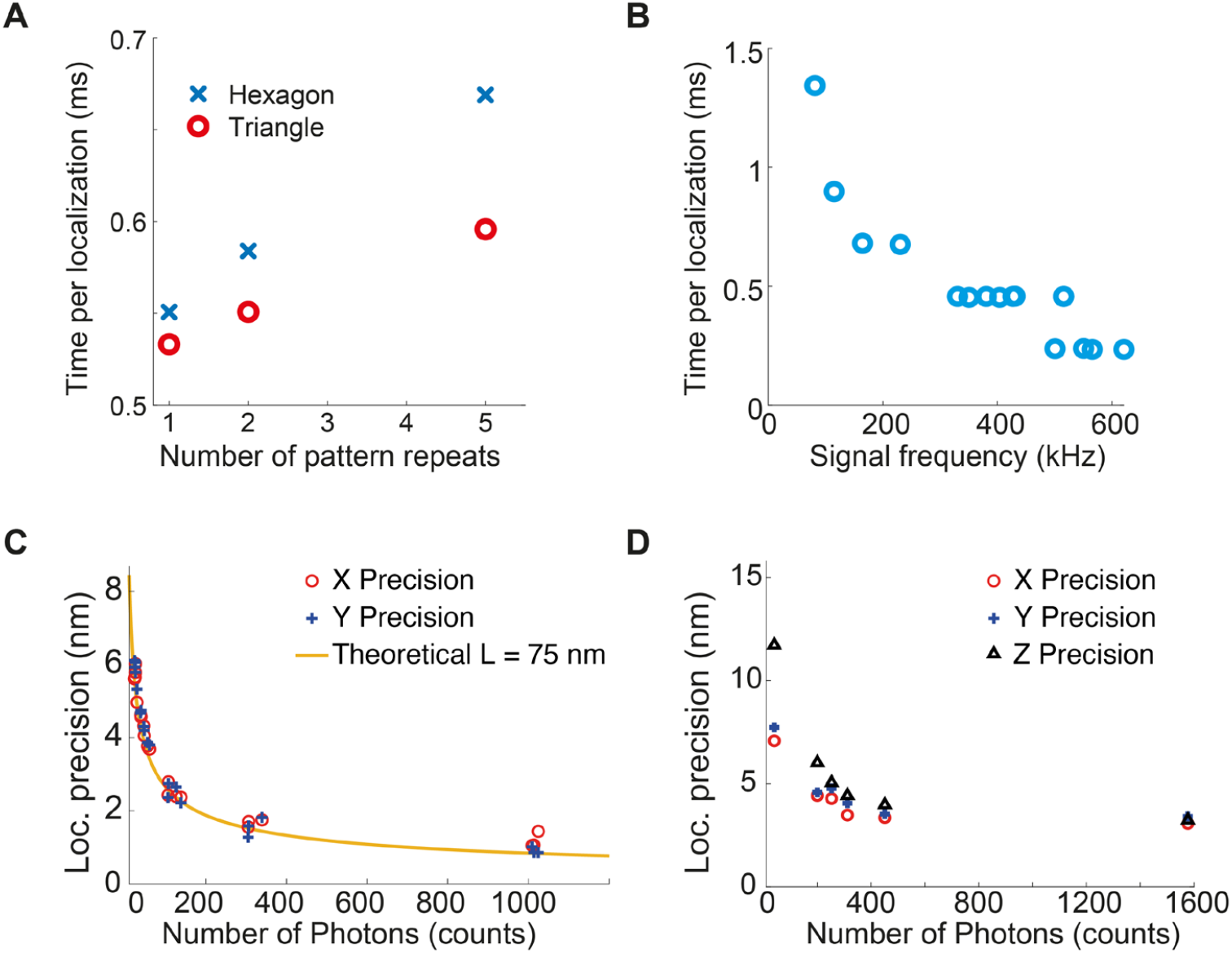
Tracking parameter optimizations. (**A - C**) **2D tracking parameter optimization**. (**A**) Single molecules of Atto647N deposited on a glass coverslip were tracked with two targeted coordinate patterns (TCP), namely hexagon and triangle, and different numbers of pattern repeats, and the temporal resolutions per localization are compared. Although the hexagonal TCP takes more time, the difference is not decisive, as our target temporal resolution is 1 ms. Due to the higher accuracy of the localization, we selected the hexagonal TCP in all our 2D tracking experiments (*14*). (**B**) Single molecules of Atto647N deposited on a glass coverslip were tracked at different laser power (thus detection signal frequency) and the temporal resolutions per localization are compared. (**C**) Single molecules Atto647N on a coverslip were tracked with different number of collected photons at the last iteration and resulting experimental localization precisions (the standard deviation of coordinate difference between consecutive localizations) are plotted. (**D**) 50 nm fluorescent nanospheres deposited on a glass coverslip were tracked in 3D and the resulting localization precisions are plotted. All the tracking parameters used in our experiments are listed in the method section.

**Fig. S2:**
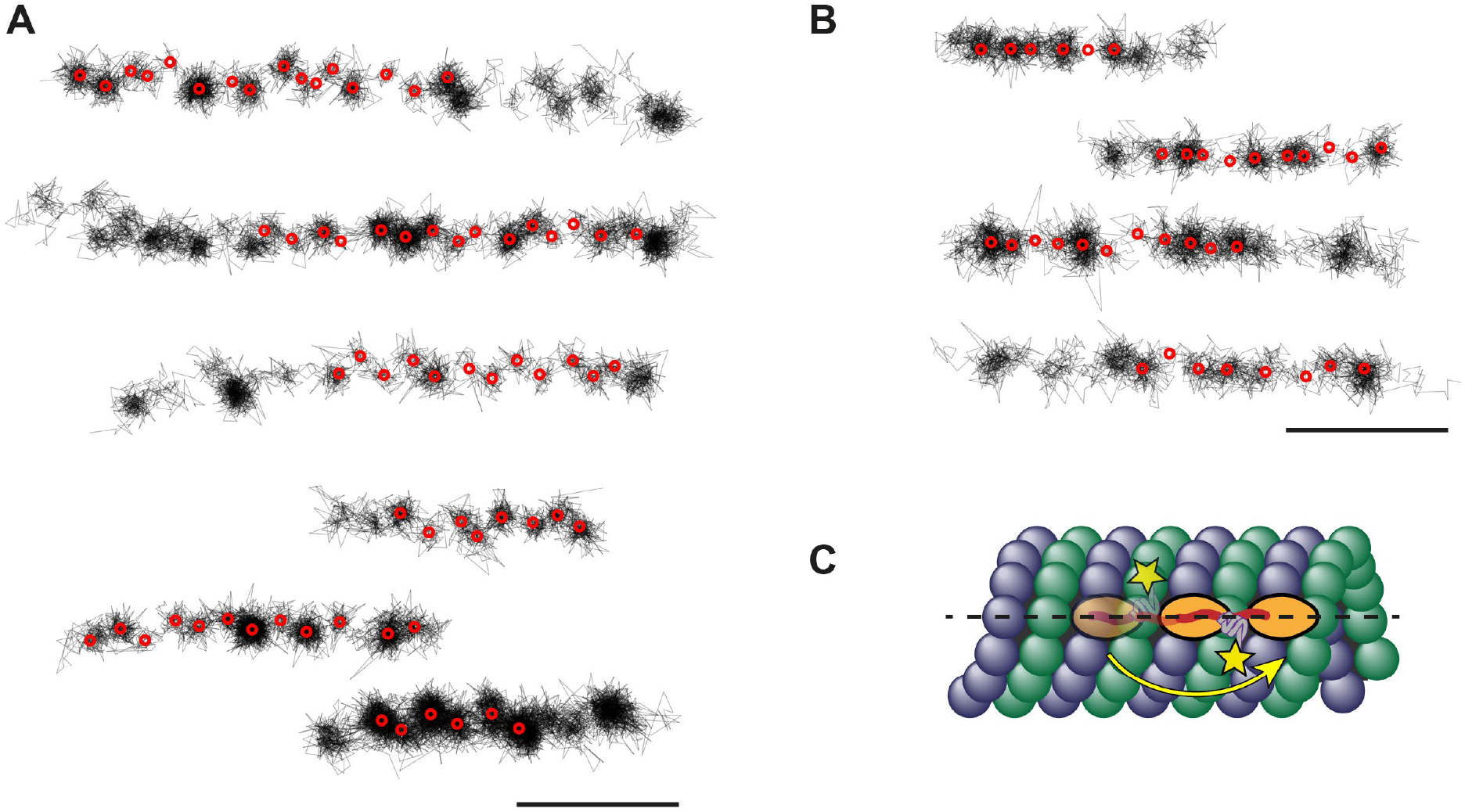
Zigzag trajectories. (**A**) Example trajectories with clear zigzag offsets. The detected steps are marked with red circles. (**B**) Example trajectories without clear zigzag offsets. (**C**) Schematic of a C-terminally labelled kinesin. When the label position is asymmetric relative to the connection of the motor domains, this will cause positional offsets perpendicular to the walking direction at each step. Scale bars: 50 nm.

**Fig. S3:**
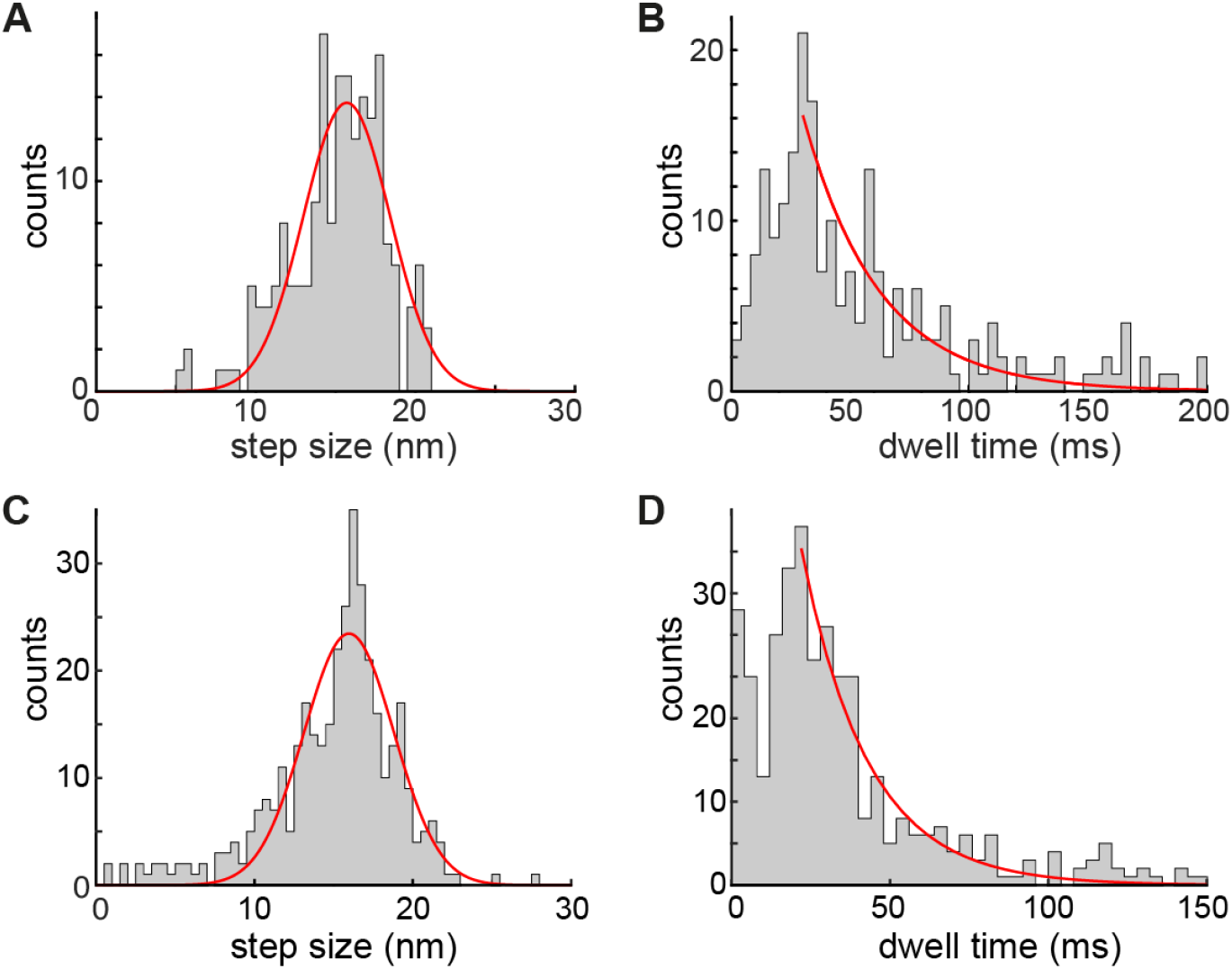
Step size and dwell time histograms. (**A, B**) Full-length wild type kinesin-1 tracking in U2OS cells. (A) A step size histogram with the Gaussian fit results in a step size of 15.7 ± 3.8 (std) ± 0.8 (sem) nm. (B) A dwell time histogram with a single exponential fit (*τ* = 31.8). Data from 6 experiments, 10 tracks, and 234 steps. (**C, D**) K560 tracking in live neurons. (C) A step size histogram with the Gaussian fit results in a step size of 16.0 ± 4.0 (std) ± 0.5 (sem) nm. (D) A dwell time histogram with a single exponential fit (*τ* = 20.9 ms). Data from 10 experiments, 25 tracks, and 409 steps.

**Fig. S4:**
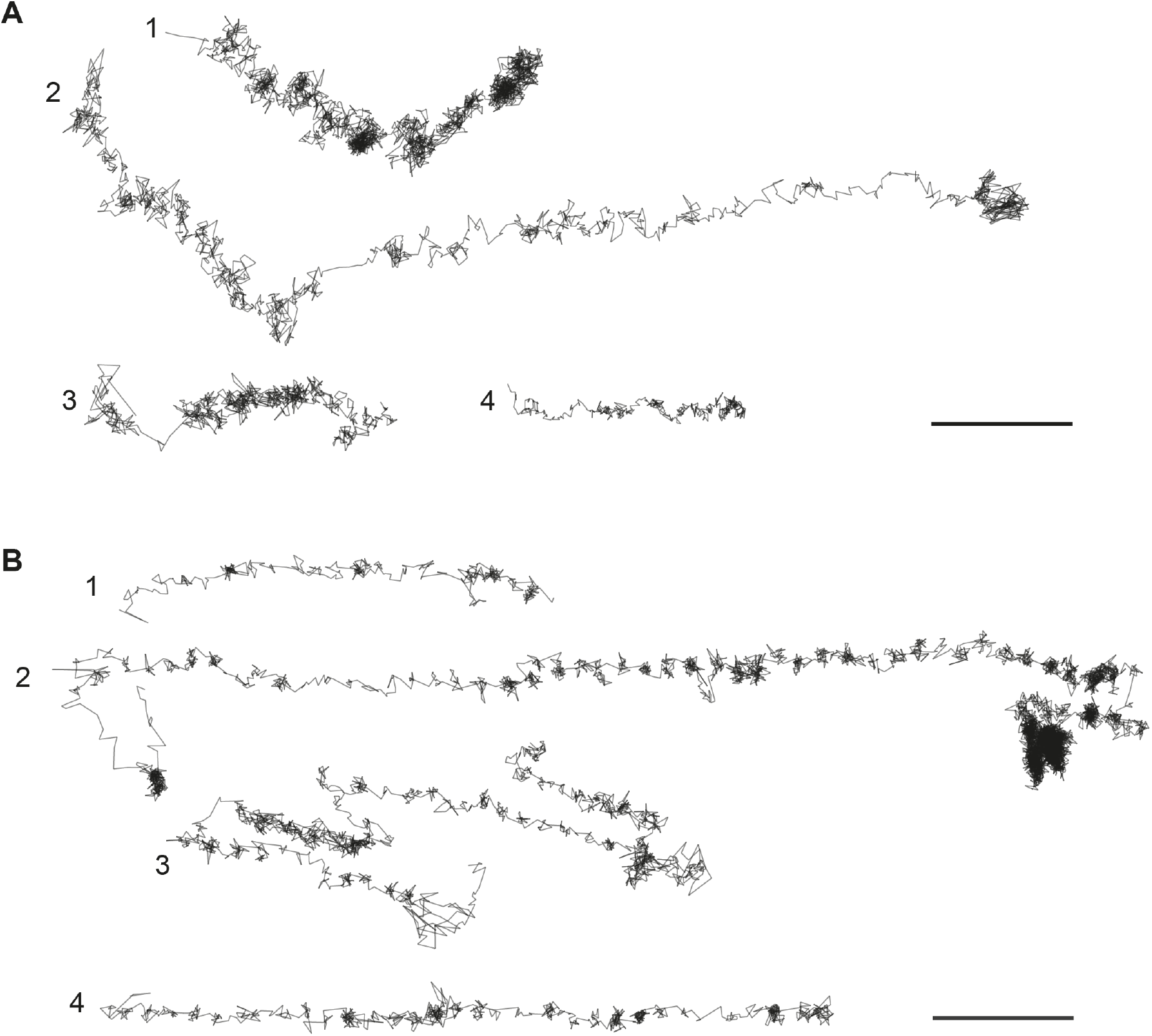
Example trajectories of full-length wild type kinesin-1. (**A**) Trajectories without clear steps, indicating passive kinesin transports by attached cargos. (**B**) Trajectories with clear steps indicating active kinesin transports. Scale bars: 100 nm.

**Fig. S5:**
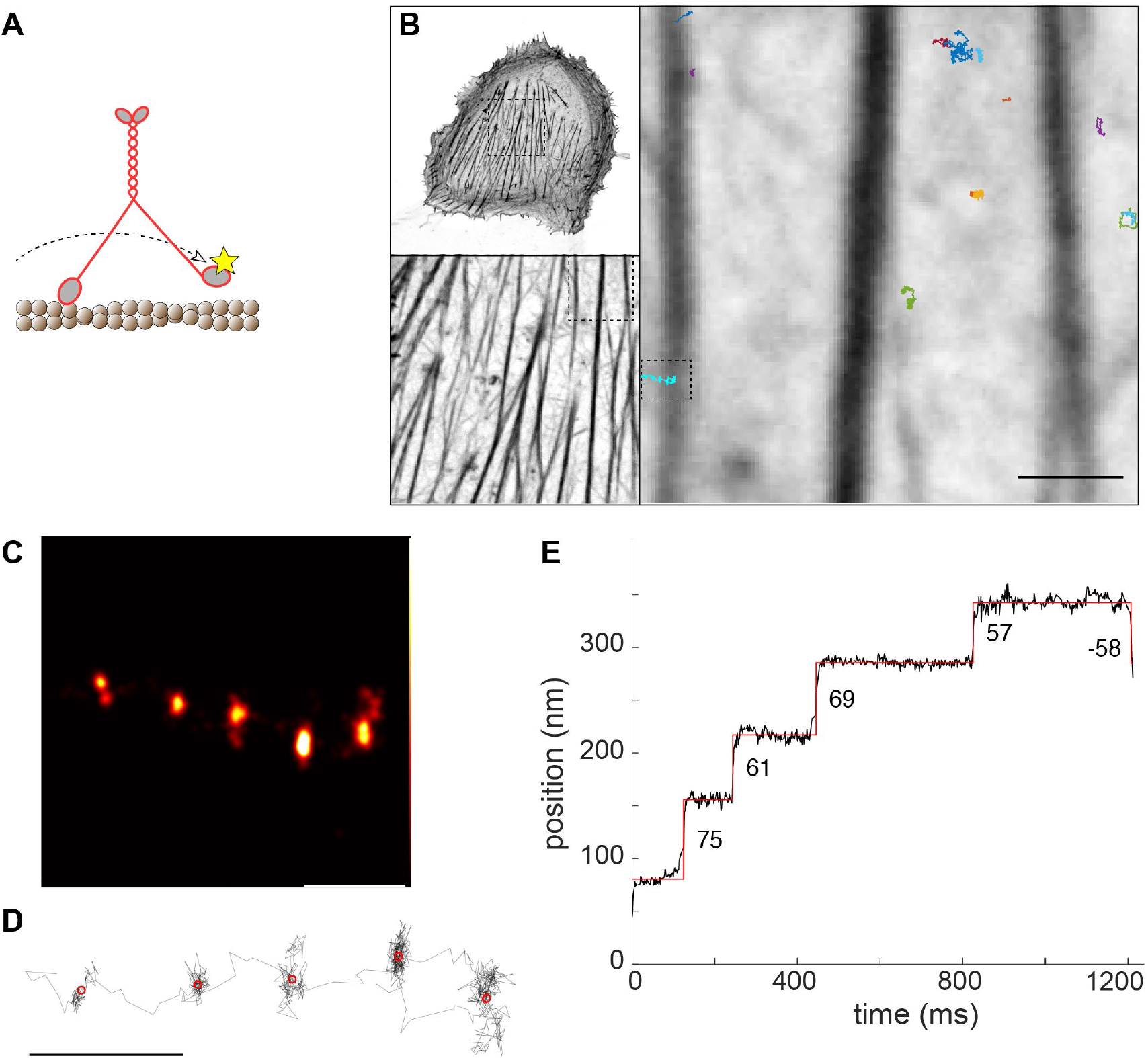
Myosin-V tracking in live U2OS cells. (**A**) Myosin-V, fluorescently labelled at the N-terminus, walks in a hand-over-hand manner with a step size of 74 nm. (**B**) Confocal images of actin-GFP in a U2OS cell, and overlaid trajectories of human myosin-V (K1701A mutant), labeled with Halo-JF646. (**C**) A myosin-V track where the localizations are rendered as a super-resolution image. (**D**) A line plot image connecting each localization. (**E**) Time versus position plot of the track, showing walking steps of around 74 nm. Scale bars: 100 nm (D), 1 µm (B).

**Fig. S6:**
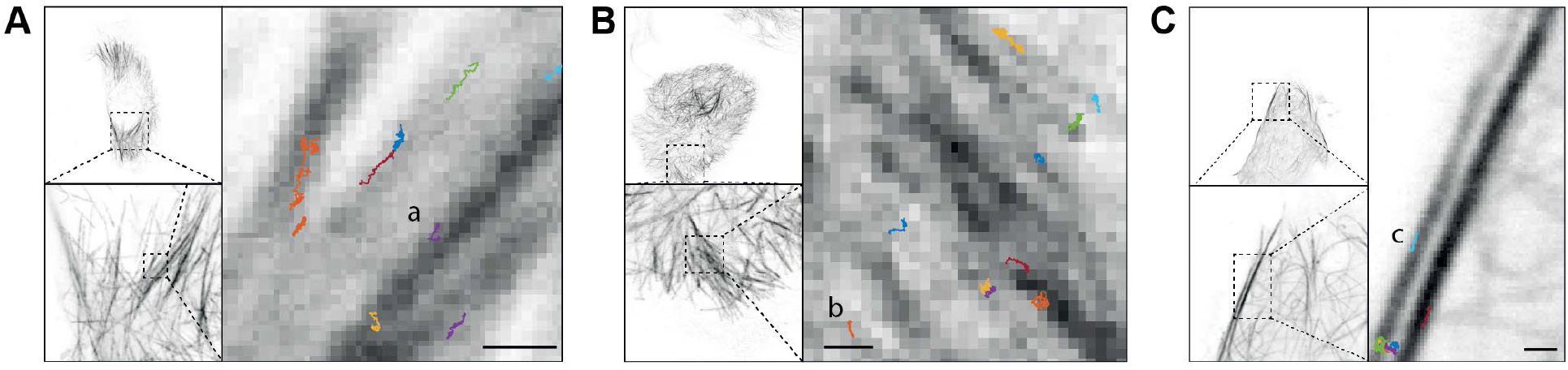
Confocal Images of live cells for 3D tracking. (**A, B, C**) Confocal images of GFP-α-tubulin in live cells where 3D tracks were captured. Tracks (a, b, c) correspond to tracks in Fig. 3 E, F. Scale bars: 500 nm (A, B, C).

**Table S1:** Experimentally obtained values of kinesin dynamics. ^1^Standard deviation from a Gaussian fit to the step size histogram. ^2^Standard error of the mean. ^3^Dwell time from fitting a single exponential function to the histogram.

|  | Velocity<br>(nm/s) | std<br>(nm/s) | Step size<br>(nm) | std<sup>1</sup> (nm) | sem<sup>2</sup><br>(nm) | Dwell<br>time<sup>3</sup><br>(ms) |
| --- | --- | --- | --- | --- | --- | --- |
| Motor-PAINT high ATP | 191 | 79 | 7.8 | 2.8 | 0.1 | 27.0 |
| FL WT kinesin-1 | 259 | 177 | 15.7 | 3.8 | 0.8 | 31.8 |
| K560 in U2OS cells | 523 | 219 | 16.2 | 3.8 | 0.06 | 19.1 |
| K560 in neurons | 381 | 214 | 16.0 | 4.0 | 0.5 | 20.9 |

## Supplementary Movies

***Supplementary movie 1***.

Kinesin (Halo-K560) motilities in a motor-PAINT U2OS cell at saturating ATP concentration, acquired by a standard fluorescence microscope. Fig.1E-H. 500 frames with exposure time of 50 ms.

***Supplementary movie 2***.

Kinesin (*Dm*KHC (1-431)-SNAP-6xHis) track in a motor-PAINT U2OS cell showing side-stepping. Fig.1G. Scale bar 10 nm.

***Supplementary movie 3***.

Kinesin (*Dm*KHC (1-431)-SNAP-6xHis) track in a motor-PAINT U2OS cell at low ATP concentration. Fig.1L. Scale bar 10 nm.

***Supplementary movie 4***.

Kinesin (*Dm*KHC (1-431)-SNAP-6xHis) track in a motor-PAINT U2OS cell at low ATP concentration showing zigzag motions. Fig.1M. Scale bar 10 nm.

***Supplementary movie 5***.

Kinesin (Halo-K560) motilities in a live U2OS cell with Taxol treatment, acquired by a standard fluorescence microscope. Fig.2E-J. 1000 frames with exposure time of 30 ms.

***Supplementary movie 6***.

Kinesin (Halo-K560) track in a live U2OS cell with Taxol treatment showing clear walking steps. Fig.2Gd. Scale bar 10 nm.

***Supplementary movie 7***.

Kinesin (Halo-K560) 3D track in a live U2OS cell with Taxol treatment acquired by 3D MINFLUX tracking. Fig.3Ec.

